# SpyTag-Enabled Assembly of Bacterial Microcompartment Trimers into Macroscopic Layered Protein Materials

**DOI:** 10.64898/2026.04.06.716716

**Authors:** Yali Wang, Xiaobing Zuo, Yaqing Wang, Paul D. Ashby, Robert P. Hausinger

## Abstract

Protein self-assembly enables precise nanoscale organization but rarely translates into macroscopic materials that retain functionality beyond aqueous environments. Here, we report that a bacterial microcompartment (BMC) trimer fused with SpyTag (T1-SpyTag), when expressed as a standalone component, undergoes rapid and spontaneous self-assembly into macroscopically visible fibers and layered sheets. These structures span from the nanoscale to the millimeter scale, forming robust three-dimensional protein materials that remain structurally intact and functionally accessible in both solution and dried states. Unlike previously reported SpyTag-enabled BMC systems that function primarily as passive cargo-loading modules, T1-SpyTag macromolecular structures exhibit emergent material behavior, including chemical robustness under denaturing conditions, while preserving covalent reactivity toward SpyCatcher-fused cargos. The multilayered architecture enables tunable surface display, access to ultrathin, processable protein films, and surface renewability through layer-by-layer removal and regeneration. This work demonstrates how a minimal genetic modification of a native protein building block can drive the formation of functional, macroscopic protein materials, thus expanding the design space of BMC-derived assemblies for biointerfaces, catalysis, and sustainable protein materials engineering.

## 1. Introduction

Protein self-assembly is a fundamental organizing principle in biology, giving rise to highly ordered structures such as viral capsids,^1^ bacterial microcompartments (BMCs),^2^ filaments,^3^ and S-layer lattices.^4^ Inspired by these natural systems, engineered proteins have been widely explored as building blocks for nanomaterials, drug delivery vehicles, biosensors, and catalytic platforms.^5-6^ Despite these advances, most reported protein assemblies remain confined to nanometer-to-micrometer length scales, typically forming nanoparticles, shells, or short filaments. Moreover, many systems are unstable outside of aqueous environments and lose structural integrity or functionality upon drying.^7^ Achieving protein-based assemblies that extend from nano to macro scales while also retaining stability and functionality in both liquid and solid states remains a major challenge in biomaterials research.

BMCs are protein-based organelles that encapsulate enzymes to optimize specific metabolic pathways.^8 9^ The shells of BMCs are constructed from three major classes of proteins: hexamers (H), trimers (T), and pentamers (P).^10^ Together, these proteins assemble into icosahedral (Fig. 1) or tubular shells, creating defined nanostructures with selective permeability.^11 12 13^Because of their inherent symmetry and modularity, BMC shell proteins have been repurposed in synthetic biology as programmable scaffolds.^14^ Among them, trimeric shell proteins fused with functional tags have been widely used to introduce additional enzymatic modules into the BMC lumen. Of special note are the covalent conjugation systems such as SpyTag/SpyCatcher that have further expanded the utility of BMC shell proteins by enabling programmable cargo loading and surface functionalization.^15^ For example, engineered BMC-T1 proteins fused with SpyTag have been employed to load fluorescent proteins or enzymes into nanoscale shells via covalent capture by SpyCatcher fusions.^15 16 17^ To date, however, such constructs have been regarded primarily as cargo-loading modules, and their intrinsic assembly behaviors have not been systematically explored.

**Figure 1.**
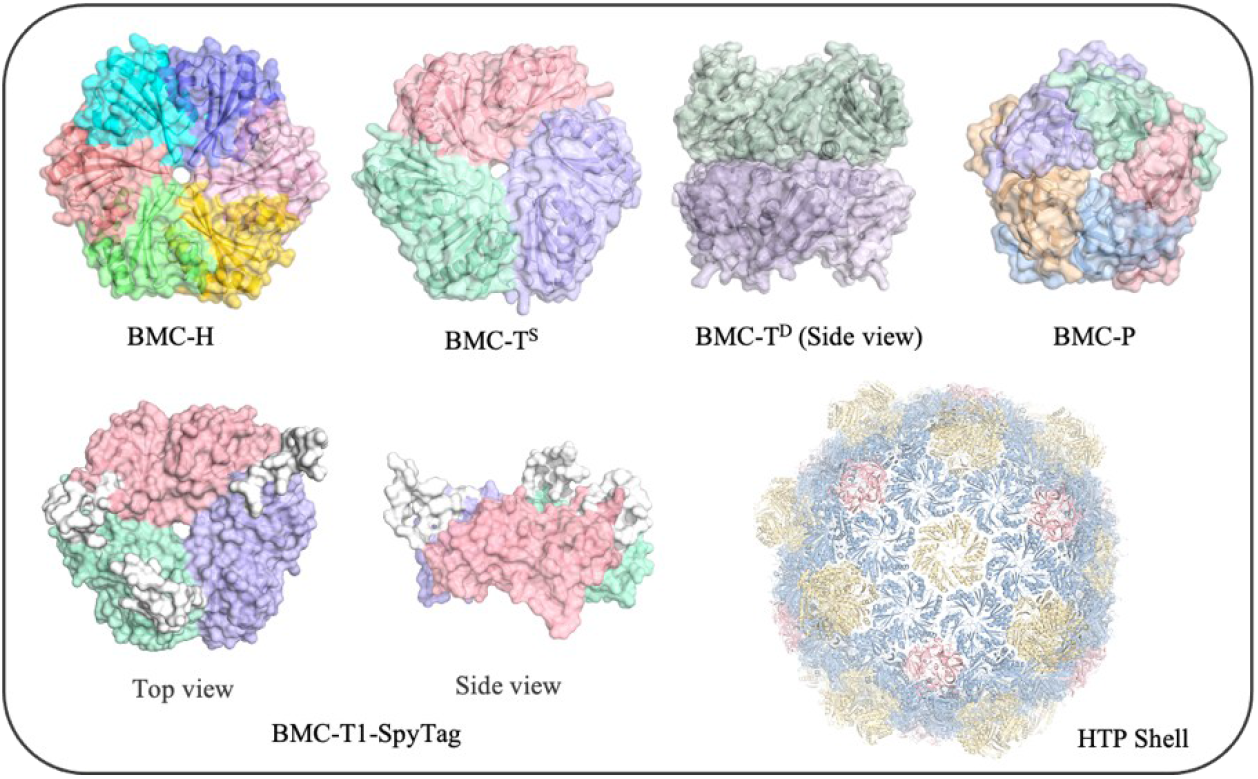
Structures of the bacterial microcompartment components, the intact HTP shell structure, and the SpyTag-fused BMC-T1 trimer. The structures of bacterial microcompartment components include the hexamer (H), single-layer T1 trimer (T^S^), double-layer trimers (T^D^) and pentamer (P), along with the assembled intact HTP shell, with hexamers (H) shown in blue, trimers in yellow, and pentamers in red. The structure of SpyTag-fused BMC-T1 trimer (T1-SpyTag) was predicted using AlphaFold, with top and side views shown and with SpyTag moieties depicted in white.

Here, we report that a BMC-T1 fused to SpyTag (denoted T1-SpyTag, Fig. 1, sequence provided in Supplementary Information) undergoes rapid and spontaneous self-assembly into macroscopically visible fibers and sheet-like materials when expressed as a standalone component. These structures reach centimeter-scale dimensions and retain covalent functional accessibility in both solution and dried states. Importantly, these assemblies exemplify how highly ordered nanoscale protein building blocks can give rise to globally disordered yet stratified materials with promising macroscopic properties, while enabling access to ultrathin, processable protein films. Our findings establish T1-SpyTag as a new class of protein material, demonstrating how a minimal genetic modification of a native protein building block can drive the emergence of functional, macroscopic structures.

## 2. Results and Discussion

### 2.1 Spontaneous rapid formation of macroscopic fibers and sheets

We repeatedly observed that the SpyTag-fused BMC-T1 (T1-SpyTag) underwent rapid self-assembly into visible fibrous and sheet-like aggregates after purification. When freshly purified protein was briefly agitated, protein fibers appeared, and when concentrated by Amicon centrifugal ultrafiltration, sheet-like protein adhered to the filter membrane surface (Fig. 2A-D). Optical microscopy confirmed that fibers extended up to ∼2.5 cm in length and sheets spanned ∼2-3 cm in width, both having defined boundaries and remaining intact after resuspension. Transmission electron microscopy (TEM) confirmed the presence of elongated fibers and flat sheet morphologies, with the fibers clearly composed of multiple bundled protein strands (Fig. S1). Importantly, the macroscopic materials were composed solely of T1-SpyTag without detectable contaminants as shown by sodium dodecylsulfate-polyacrylamide gel electrophoresis (SDS-PAGE) analysis (Fig. 2E).

**Figure 2.**
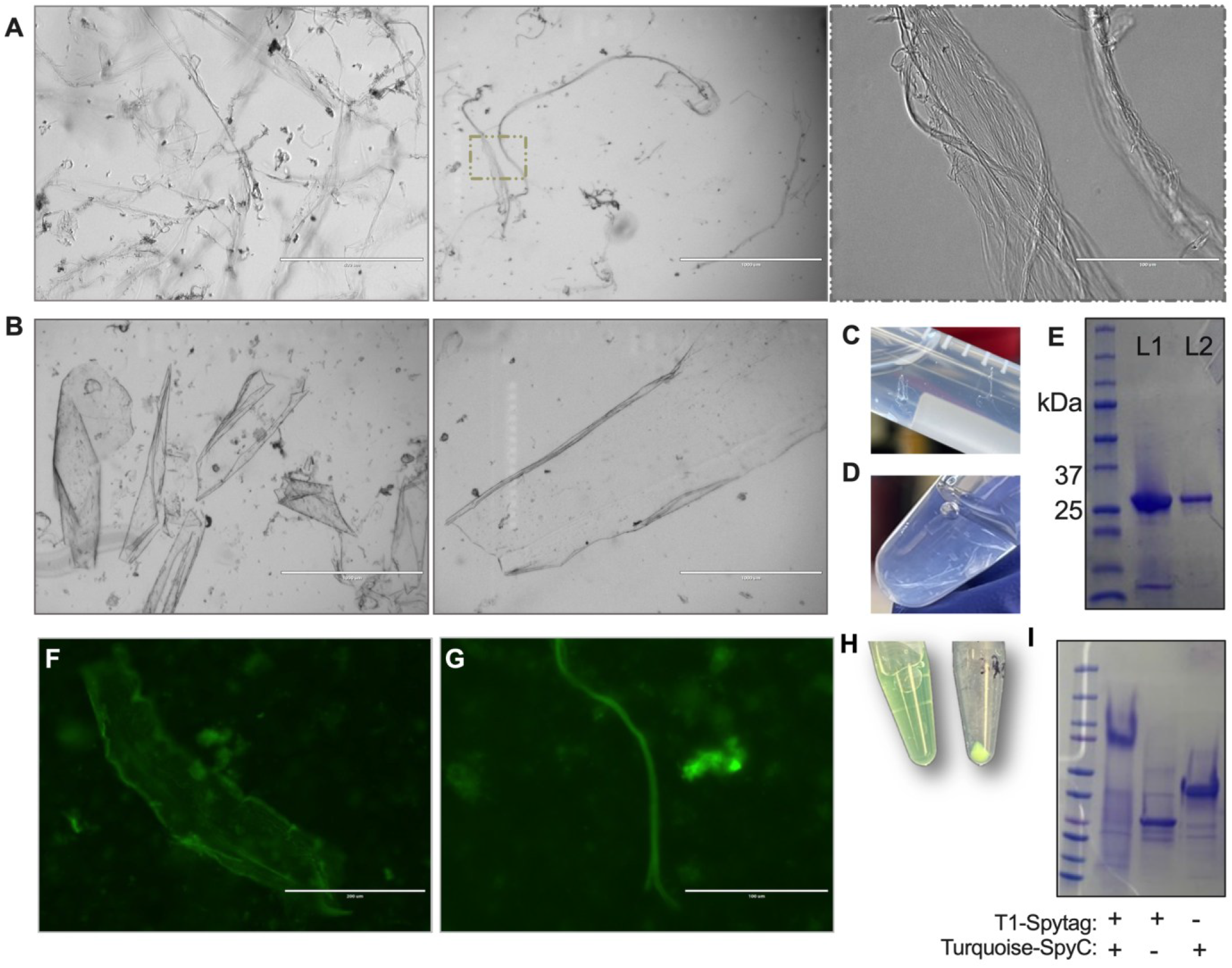
Self-assembly of T1-SpyTag into macroscopic fibers and sheets and its ability to conjugate with Turquoise-SpyCatcher (Turquoise-SpyC). Transmitted-light microscopy images of assembled T1-SpyTag showing **(A)** fibrous structures and **(B)** sheet-like structures captured using an EVOS optical microscope. In **(A)**, the right panel is an expanded view of the boxed portion of the middle panel. **(C)** Photographs of fibrous assemblies and **(D)** sheet-like assemblies formed in storage tubes. **(E)** SDS–PAGE analysis of soluble T1-SpyTag (L1) and an assembly dissociated by urea treatment (L2). Representative fluorescence microscopy images demonstrating the conjugation of Turquoise-SpyC with T1-SpyTag assemblies for **(F)** sheets and **(G)** fibers. **(H)** Photograph of the Turquoise-SpyC–T1-SpyTag conjugation samples in solution and after centrifuging. **(I)** SDS–PAGE analysis of the conjugate of a T1-SpyTag assembly and Turquoise-SpyC, along with the non-conjugated components.

Previous studies have shown that BMC shell proteins are evolutionarily optimized for modular assembly and selective permeability, and are classically known to form nanoscale cages and curved shell structures associated with intracellular compartments (Fig.1).^18^ BMC hexamers can assemble into ordered structures at the nano-to submicron scale,^13 19^ or they can form patterned assemblies at liquid–liquid or liquid–solid interfaces.^20^ In contrast, the rapid emergence of centimeter-scale fibers and sheets observed here demonstrates that a BMC-derived trimer can intrinsically access stable, three-dimensional macroscopic assemblies, a behavior not previously reported for other BMC H/T/P shell proteins.

### 2.2 T1-SpyTag assemblies preserve full bioactivity in solution

SpyTag/SpyCatcher has been widely used in a modular strategy for cargo loading and surface functionalization in protein assemblies,^21 22 23 5^ including engineered BMC shells, where SpyTag serves primarily as a passive conjugation handle for post-assembly attachment of functional cargos.^14 16^ A recent study further demonstrated that the BMC shell components H, T, T-SpyTag, and P, when independently expressed and purified, can assemble *in vitro* into intact shells and enable cargo loading via SpyCatcher fusion.^2^

To determine whether the T1-SpyTag supramolecular assembly alters the accessibility of the SpyTag component, pre-formed T1-SpyTag sheets and fibers were incubated with SpyCatcher– Turquoise fusion protein (abbreviated Turquoise-SpyC, with the sequence provided in Supplementary Information). Following extensive washing, strong and specific turquoise fluorescence was detected throughout both architectures (Fig. 2F-H), whereas no signal appeared in the absence of Turquoise-SpyC. The uniform fluorescence distribution across both sheets and fibers indicated that the SpyTag moieties remain exposed on the surfaces of the assemblies and that the structures maintain mesoscale order despite their macroscopic size. This functional persistence after assembly contrasts sharply with many protein materials, where domain burial and loss of reactivity are typical. SDS–PAGE analysis of urea-dissolved assemblies revealed a single band corresponding to the covalent T1-SpyTag–Turquoise-SpyC conjugate (Fig. 2I), confirming the formation of an intact isopeptide bond. Thus, self-assembly does not occlude or deactivate SpyTag residues on the solvent-accessed surface, even after assembly into micron-scale to centimeter-scale architectures. This finding establishes that minimal genetic modification can simultaneously retain molecular recognition functionality and drive BMC-derived building blocks into a macroscopic materials regime.

### 2.3 Concentration-dependent initiation and kinetics of assembly

The protein concentration exerted a strong effect on the kinetics of T1-SpyTag self-assembly in solution. To assess this effect, varied concentrations of the protein were incubated under static conditions and spectra were recorded from 300–450 nm to monitor light scattering. After 10 min of incubation, the samples were essentially indistinguishable across concentrations, consistent with no differences in the amount of scattering and thus no measurable assembly at these early times (Fig. 3A). After 24 h of undisturbed incubation, the readouts diverged markedly (Fig. 3B) with solutions at 13 μM or less remaining optically clear and showing negligible spectral change compared to the TBS buffer control, and with samples at concentrations of 20 μM and greater exhibiting strong wavelength-dependent baselines consistent with enhanced light scattering. Furthermore, small sheet-like structures became clearly visible by eye in the wells. Intermediate concentrations produced graded increases in the scattering baseline, consistent with a concentration-driven amplification of assembly. Notably, increasing the T1-SpyTag concentration beyond ∼40 μM resulted in little further enhancement of either the rate or the extent of scattering, consistent with an apparent critical concentration above which the self-assembly process approaches saturation. By monitoring the absorbance at 380 nm over 15 h, we revealed that, under static conditions, assembly differences between groups began to emerge after approximately 9.5 hours (Fig. 3C).

**Figure 3.**
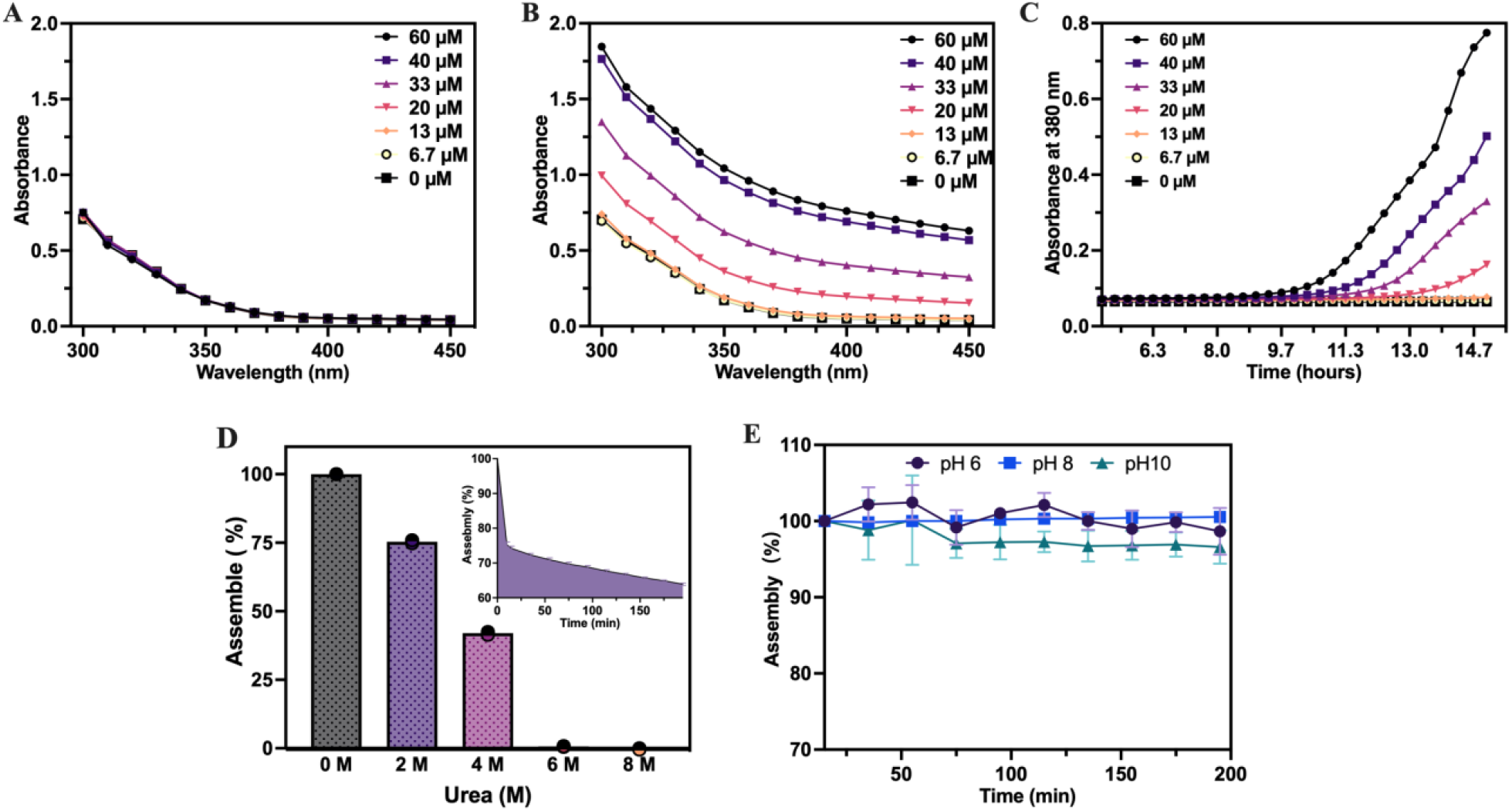
Assembly and disassembly behavior of T1-SpyTag macromolecular structures. UV– visible absorption spectra (300–450 nm) of T1-SpyTag in solution were recorded at room temperature after **(A)** 10 min and **(B)** 24 h using varied protein concentrations, with TBS as the control. The apparent absorbance changes reflect differences in light scattering due to protein assembly into macromolecular structures. **(C)** Time-dependent changes in absorbance at 380 nm during the assembly of T1-SpyTag over 15 h. **(D)** Disassembly of T1-SpyTag structures when incubated with different urea concentrations. The inset shows the time-dependent decrease of the assembly ratio in the presence of 2 M urea. **(E)** Stability of T1-SpyTag macroscopic structures at different pH values of 6, 8, and 10.

Together, these results demonstrate that T1-SpyTag assembly in static conditions is governed by a critical concentration threshold. Once this threshold is exceeded, assembly proceeds cooperatively and leads to rapid formation of macroscopic structures, whereas below the threshold nucleation is strongly suppressed. This finding explains our initial observations: the pronounced increase in T1-SpyTag concentration when using an Amicon membrane led to the rapid generation of centimeter-scale sheet structures. This concentration-dependent behavior underscores the intrinsic capacity of T1-SpyTag to undergo spontaneous, threshold-driven self-assembly under both static and dynamic conditions.

### 2.4 Chemical robustness of T1-SpyTag macromolecular structures

After air-drying the macromolecular T1-SpyTag structures in 96-well plates, we observed smooth, white protein films that adhered tightly to the bottom surface and exhibited a layered morphology. To evaluate the robustness of these dried assemblies, we subjected the T1-SpyTag biomaterial to urea treatment and to TBS buffers of varying pH while monitoring the apparent absorbance at 350 nm.

Notably, the dried T1-SpyTag sheets maintained ∼75% and ∼42% integrity after 10 min in 2 M and 4 M urea, respectively, as judged by the changes in light scattering (Fig. 3D). In 2 M urea, the disassembly kinetics followed a distinct two-stage pattern, consisting of a rapid initial phase within the first 10 min, followed by a significantly slower dissociation process (Fig. 3D, inset). Remarkably, even after 2 h of 2 M urea challenge, ∼63% of the assembled structure remained intact, underscoring the extraordinary resilience of the material. This behavior is consistent with a multilayer arrangement, where the internal layers exhibit greater density and stability, while the outer surfaces are relatively more accessible and therefore more susceptible to disassembly. Significantly, the dried T1-SpyTag sheets remained highly stable across a broad pH range (pH 6–10), with negligible changes in assembly ratio over 200 min, indicating that the material retains its structural integrity under both mildly acidic and alkaline conditions (Fig. 3E).

Among self-assembling protein materials, few systems maintain structural integrity when incubated with more than 2 M urea without covalent crosslinking. The ability of T1-Spytag to withstand 2–4 M urea and the wide pH range further distinguish it as a mechanically and chemically robust protein-based material.

### 2.5 Sheet architecture

Scanning electron microscopy (SEM) imaging revealed that dried T1-SpyTag assemblies form continuous free-standing sheet-like films with micron-scale thickness. Low-magnification top-view images showed that the upper surfaces of the films are remarkably flat and featureless over large lateral areas, indicating the formation of an interfacial-driven flattened and densely packed outer layer during solvent evaporation (Fig. 4A). Higher magnification images revealed fractured cross-sections that exposed the internal stratified architecture of the films (Fig. 4B). Some fractured regions displayed clean, planar cleavage surfaces, consistent with fracture occurring along interlayer boundaries. In contrast, other regions showed rough, jagged, and torn morphologies, with visible fibrillar textures bridging adjacent layers. This heterogeneous fracture behavior indicates that the T1-SpyTag sheets are likely composed of vertically stacked lamellar domains with non-uniform interlayer cohesion. Corresponding side-view SEM images (Fig. 4A) directly measured film thicknesses in the range of approximately ∼7 μm, confirming the macroscopic integrity of the sheets.

**Figure 4.**
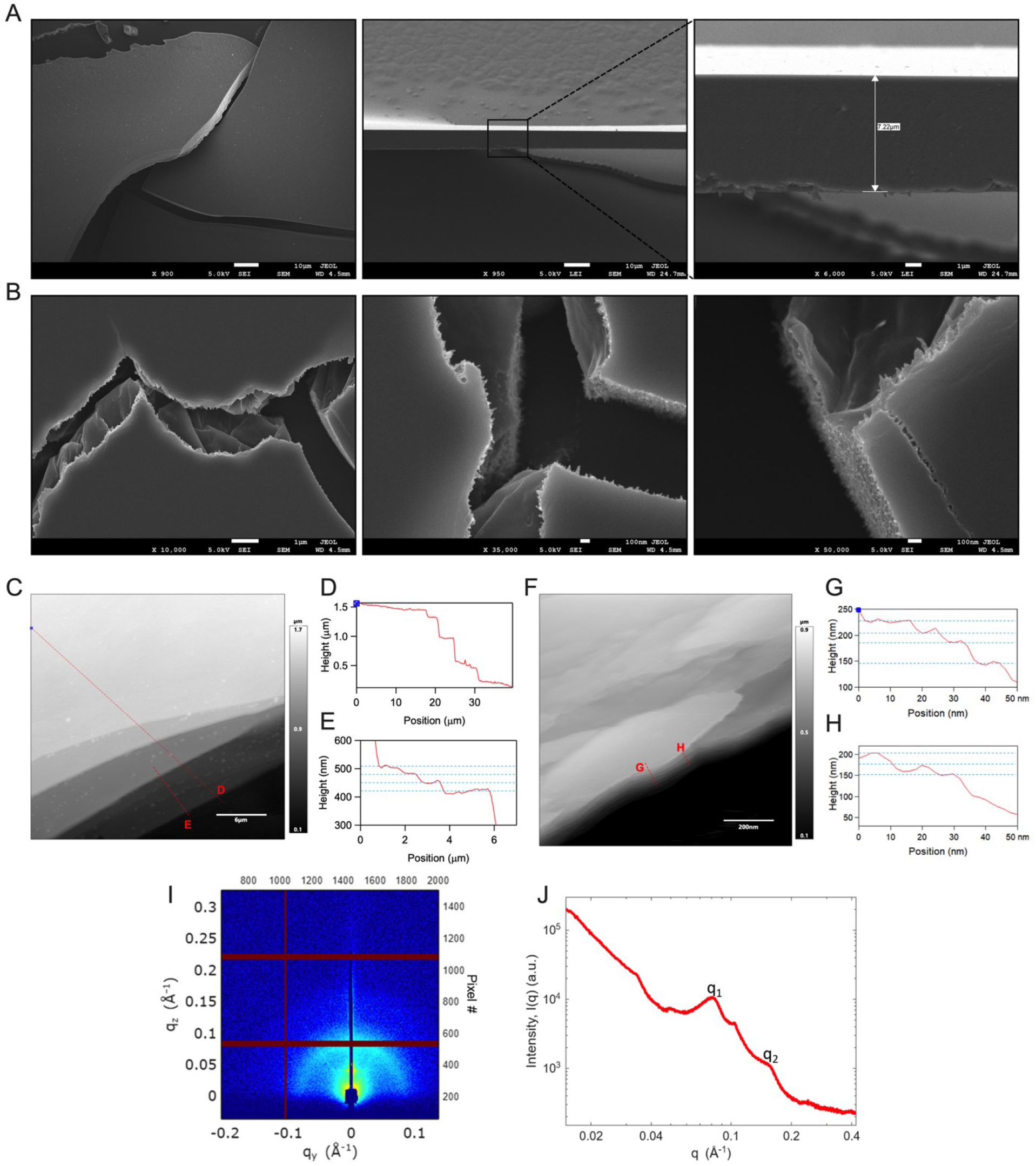
Morphology and cross-sectional architecture of self-assembled dry T1-SpyTag protein sheets. **(A)** Representative SEM images of evaporated, self-assembled T1-SpyTag sheets showing smooth upper surfaces and well-defined cross-sections with an average thickness of approximately 7 μm. **(B)** High-magnification SEM images revealing fracture features and densely stacked internal layered structures. **(C)** AFM height image of an air-dried T1–SpyTag sheet. The red lines indicate the positions used for height profiling in **(D)** and **(E)**, revealing nano-to micron-scale stepwise height transitions between stacked lamellae. **(F)** AFM height image of T1–SpyTag sheets deposited from TBS buffer. **(G)** and **(H)** Corresponding cross-section height profiles measured along the indicated lines in **(F). (I)** 2-D grazing-incidence transmission small angle X-ray scattering (GTSAXS) image for the T1-SpyTag sheet sample. **(J)** 1D SAXS profile converted from the GTSAXS image data in **(I)**. There are two major diffraction peaks observed in the 1D SAXS profile, labeled as q_1_ and q_2_. q_1_=0.079±0.009 Å^-1^, and q_2_=2*q_1_.

To examine whether the nanoscale assembly structural is conserved under different macroscopic formation routes such as air-drying versus membrane-induced concentration, we compared the atomic force microscopy (AFM) height profiles of air-dried films and preformed sheets in solution. The air-dried sample exhibited clear stepwise height transitions across stacked lamellae spanning nano-to micron-scale thicknesses (Fig. 4C,D). In the region shown in Fig. 4E, individual step heights were typically ∼30 nm, and thicker areas increased in discrete multiples of this value, indicating that film growth proceeds predominantly through layer-by-layer stacking rather than uniform compaction.

For the preformed sheets obtained during membrane concentration in solution (Fig. 4F–H), the minimal stepwise height between adjacent lamellae was likewise on the order of ∼30 nm. The measured thickness in Fig. 4G-H variations (≈120–150 nm) correspond to approximately 4–5 stacked lamellar of 30 nm. These observations suggest that T1–SpyTag adopts a consistent basic-height and multilamellar stacking mechanism and regardless of whether assembly is driven by evaporation or confinement-induced concentration.

Additional support for a layered architecture was obtained by grazing-incidence transmission small-angle X-ray scattering (GTSAXS). After transferring a preformed T1–SpyTag sheet onto a silicon wafer, two major diffraction peaks were noted. The first peak was observed at q_1_ = 0.079 ± 0.009/Å, corresponding to a d-spacing of 2π/q_1_ = 80 ± 9 Å. The second peak appeared at q_2_ = 2*q_1_, indicating that T1-SpyTag forms a laminar, layered structure (Fig. 4G-H). The interlayer distance in the most fundamental unit is approximately 80 Å.

Combining these lines of evidence, we propose that T1–SpyTag trimers first associate in a back-to-back manner, analogous to the T^D^ oligomeric state observed for native BMC-T building blocks (Fig. 1), while preserving SpyTag exposure on the accessible surface. In this model, a minimal double-layer stacking of this unit would yield a characteristic thickness of ∼15 nm, consistent with thinner structural features detected in preformed sheets by AFM (Fig. S2). Further stacking of such double-layer modules would produce an interlayer distance on the order of ∼80 Å, in agreement with the lamellar periodicity extracted from GTSAXS measurements. Notably, the predominant step height observed by AFM is ∼30 nm, indicating that the assemblies most commonly adopt a lamellar unit consisting of two stacked ∼15 nm double-layer modules corresponding to a four-layer trimer arrangement, which may represent a particularly stable structural state within the multilayered material.

Overall, these combined SEM, AFM, and GTSAXS analyses support a hierarchical multilamellar architecture in which T1–SpyTag assembles through discrete stacking of nanoscale lamellar modules. Importantly, the conserved ∼30 nm step height observed under both air-drying and membrane-induced concentration indicates that the same fundamental assembly principle governs sheet formation across distinct macroscopic routes. This modular stacking behavior provides a structural basis for the exceptional robustness and processability of the macroscopic T1–SpyTag films.

### 2.6 Cargo-loading functionality of dehydrated T1-SpyTag

Building on our findings that the SpyTag residues are exposed and functional on T1-SpyTag assemblies in solution, and that dried T1-SpyTag films exhibit the same stepped lamellar architecture, we next examined whether this well-defined solid-state surface also preserves biochemical functionality after dehydration. Evaporated T1-SpyTag films (Fig. 5B) were incubated with Turquoise-SpyC and subjected to extensive washing to remove unbound protein. Upon dissolution of the deposited films and SDS-PAGE analysis (Fig. 5C), two distinct bands were observed corresponding to the T1-SpyTag–Turquoise-SpyC conjugate and unconjugated T1-SpyTag. Strong and spatially confined fluorescence was retained exclusively on the film surface (Fig. 5A). Combined, these results indicate that the outermost lamellae remain solvent-accessible and participate in covalent coupling, whereas inner layers remain unreacted. The homogeneous fluorescence observed across both flat terraces and exposed edges is fully consistent with the layer-by-layer stacked architecture resolved by AFM and SEM, confirming that functional SpyTag residues are continuously presented across successive sheet layers—even in the dry state.

**Figure 5.**
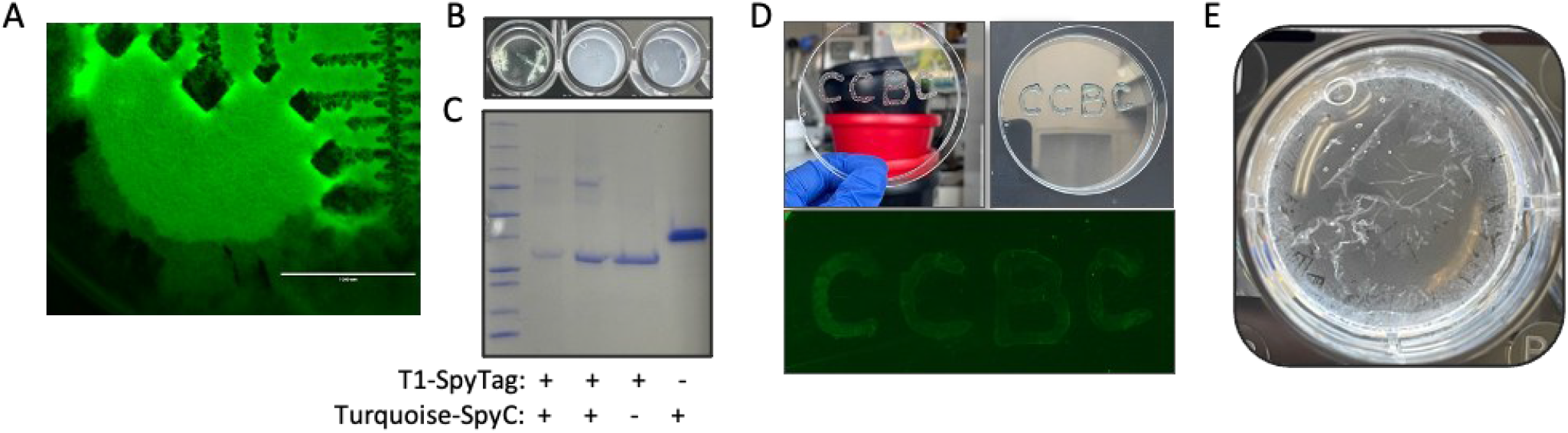
Dry-state functionality of T1-SpyTag assemblies and formation of ultra-thin sheets. **(A)** Fluorescence microscopy image of air-dried T1-SpyTag films after incubation with Turquoise-SpyC, showing strong turquoise fluorescence selectively localized to the film regions. **(B)** Representative photographs of evaporation-derived T1-SpyTag films in 96-well plates before (right two wells) and after conjugation with Turquoise-SpyC (left well). **(C)** SDS–PAGE analysis of dissolved samples of dry T1-SpyTag films and their conjugates with Turquoise-SpyC. Lanes 1–2: conjugated T1-SpyTag–Turquoise-SpyC complex (68.0 kDa); lane 3: T1-SpyTag control (25.3 kDa); lane 4: free Turquoise-SpyC (42.7 kDa). **(D)** Patterned display of T1-SpyTag films. Letter patterns (“CCBC”) were written using T1-SpyTag solution and air-dried to form visible films (top left). After incubation with Turquoise-SpyC (top right), a close-up view by fluorescence microscopy revealed turquoise signals specifically confined to the patterned regions after washing (bottom). **(E)** Photographs of ultra-thin T1-SpyTag sheets isolated from dried T1-SpyTag films.

### 2.7 Display and isolation of ultrathin T1-SpyTag sheets

The robust dry-state reactivity of T1-SpyTag films, together with their laminated architecture, enables user-defined spatial activation. Letter-shaped deposits of T1-SpyTag produced sharply resolved fluorescent “CCBC” patterns after labeling with Turquoise-SpyC (Fig. 5D), demonstrating that continuous stacked sheets remain uniformly conjugation-competent at the exposed outer lamellae. These results establish dried T1-SpyTag films as programmable, solid-state protein biointerfaces capable of mutli-scale patterning.

By tuning the ratio of deposited protein to surface area, the thickness and degree of lamellation in evaporation-assembled T1-SpyTag films can be systematically controlled. When 2 mg of T1-SpyTag was dried in a 12-well plate (surface area ∼3.8 cm^2^), the entire well bottom became coated with a thick multilayered film. Upon gentle rinsing with water, the upper lamellae delaminated cleanly and floated off as ultrathin sheets (Fig. 5E), whereas a tightly adhered basal layer remained anchored to the substrate. Repeated rinsing released successive ultrathin sheets until only a final solid layer could no longer be exfoliated. SEM images show surface cracks and wide interlayer gaps in near-surface regions, suggesting a laminated architecture that may permit layer separation (Fig. S3). Importantly, the stepwise delamination behavior is strikingly consistent with the two-phase dissociation kinetics observed for T1-SpyTag in 2 M urea.

Therefore, a defining feature of the macroscopic T1-SpyTag structures is their unusual combination of chemical robustness and retained functional accessibility.^24^ Unlike many protein-based materials that either dissociate under denaturing conditions or lose surface reactivity upon assembly,^25 26 27 28^ T1-SpyTag materials remain partially intact at high urea concentrations while preserving accessible SpyTag functionality in solution and after drying and painting. These properties arise from a multilayered architecture, in which stacked protein layers collectively provide structural integrity while maintaining reactive outer surfaces. Such assemblies may represent kinetically stabilized material states, in which structural integrity is maintained across hydrated and dry conditions without requiring irreversible densification. This organization enables tunable surface display through control of protein loading and exposed area, as well as the extraction of ultrathin protein films from surface layers without disrupting the underlying bulk material (Fig. 5E). Such layer-by-layer accessibility further suggests a route toward reusable functional materials, where surface-bound cargos can be refreshed through removal and regeneration of outer layers rather than complete material reconstruction. This layer-by-layer accessibility further opens opportunities for iterative or sequential functionalization, enabling dynamic biointerfaces with temporally controlled surface composition.

Overall, structural precedents support the feasibility of T1-Spytag assembly. High-speed atomic force microscopy has directly visualized facet-like, two-dimensional assemblies formed by BMC hexamers, revealing lateral interactions central to shell formation.^19^ Given the hexagonal structural similarity between BMC trimers and hexamers at planar interfaces, and their shared participation in native shell assembly, these observations provide a basis for the lateral association of trimers (Fig. 1 and 6A).^29 30^ Specifically, large-scale macroscopic fibers and sheets were not observed during purification of standalone T1. Thus, SpyTag extends these interactions beyond planar facets into a three-dimensional architecture (Fig. 6B), producing macroscopically visible fibers (Fig. 6C) and thick sheets (Fig. 6D) in solution rather than remaining confined to nanoscale 2D assemblies. The coexistence of fiber- and sheet-like morphologies indicates that T1-SpyTag assemblies can adopt distinct material states under different experimental contexts, including multilayered assemblies in air-dried samples (Fig. 6E-F). Future work will explore how genetic and architectural parameters of BMC-derived building blocks can be leveraged to program material dimensions and morphology.

**Figure 6.**
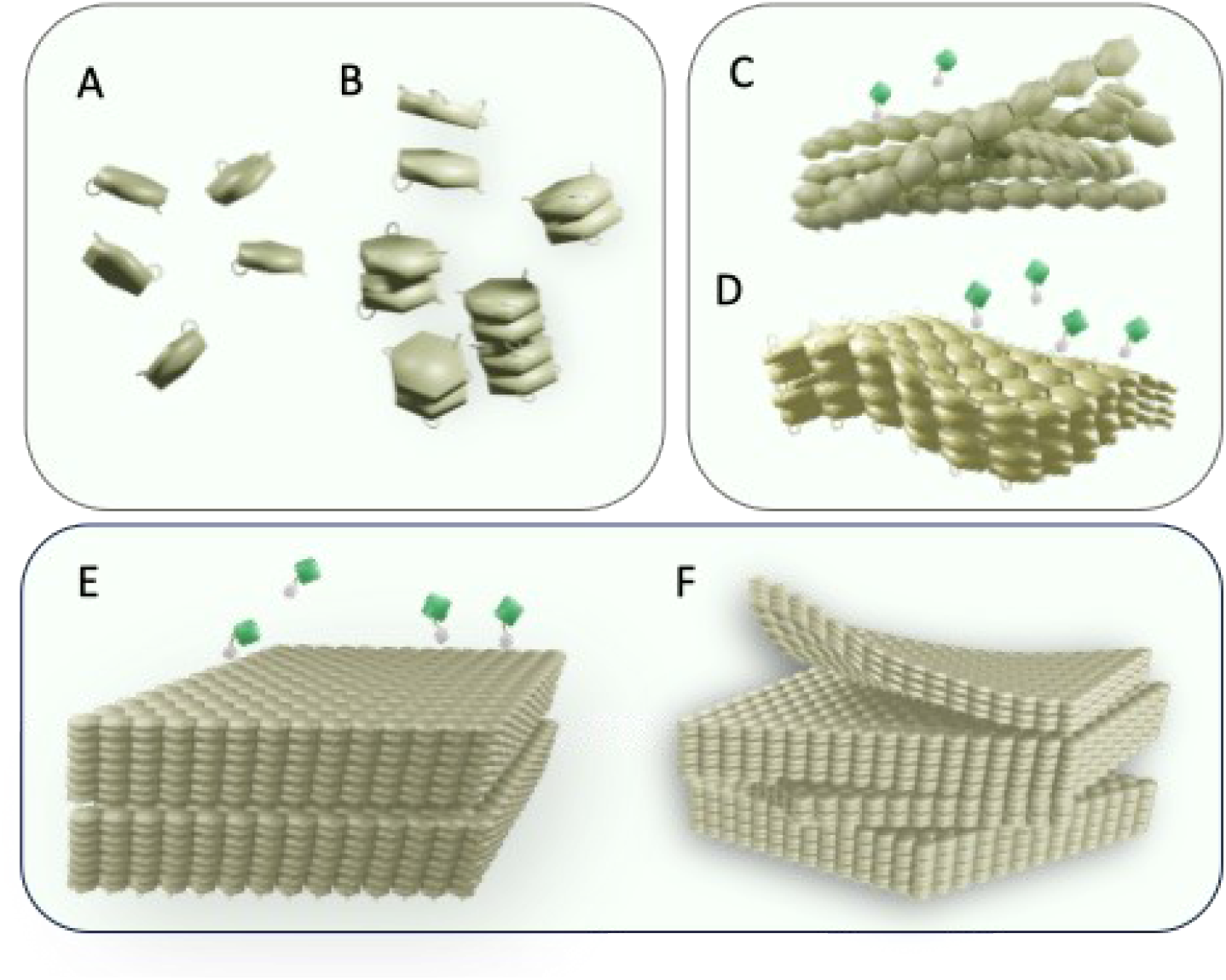
SpyTag-enabled assembly of BMC T1 trimers into macroscopic, layered protein materials. **(A)** Individual T1-SpyTag units are depicted as discrete building blocks in solution. **(B)** Local association of T1–SpyTag trimers gives rise to initial stacked assemblies depending on local concentration and surface or confinement conditions. **(C**,**D)** T1–SpyTag assemblies evolve into distinct macroscopic morphologies, including fiber-like bundles **(C)** and extended sheet-like structures **(D)** in solution, while SpyTag motifs remain exposed on accessible surfaces and able to react with SpyCatcher fusions such as Turquoise-SpyC (green and white symbols). **(E)** Multilayered assemblies of air-dried T1–SpyTag with accessible outer surfaces. **(F)** The layered architecture enables selective removal of outer layers to generate ultrathin protein films while preserving the underlying core material.

## 3. Conclusion

Collectively, this work establishes a new paradigm for BMC-derived protein materials. By minimally modifying a native BMC trimer with SpyTag, we demonstrate that building blocks traditionally associated with nanoscale biological compartments can be reprogrammed into macroscopic, three-dimensional materials that integrate chemical robustness, functional accessibility, and surface renewability. More broadly, this study highlights a general strategy for transforming genetically encoded protein assemblies into adaptive, reusable bio-based materials, expanding the design space for protein materials across biointerfaces, catalysis, and sustainable materials engineering.

## 4. Experimental Section

### Protein expression and purification

The T1-SpyTag construct was expressed in *Escherichia coli* BL21 (DE3) cells that were grown in LB medium at 37 °C to an OD_600_ of 0.6– 0.8, followed by induction with anhydrotetracycline (100 ng mL^−1^ final concentration) and incubation at 25 °C overnight. Cells were harvested by centrifugation (8,000 × *g*, 15 min) and resuspended in lysis buffer (20 mM Tris–HCl, 250 mM NaCl, 20 mM imidazole, pH 7.6). Cell disruption was performed using a French pressure cell, and the lysate was clarified by centrifugation (38,000 × *g*, 60 min). The supernatant, which contained soluble T1-SpyTag, was loaded onto a TALON® metal affinity resin (TaKaRa) gravity column, washed with lysis buffer, and treated with elution buffer (20 mM Tris–HCl, 250 mM NaCl, 300 mM imidazole, pH 7.6). Eluted proteins were concentrated using Amicon Ultra centrifugal filters (50 kDa cutoff) and buffer-exchanged into TBS buffer (20 mM Tris–HCl, 250 mM NaCl, pH 7.6) as needed for downstream experiments.

The SpyCatcher–Turquoise fusion protein (abbreviated Turquoise-SpyC, sequence provided in Supplementary Information) was obtained from a reported previously construct.^14 31^ Expression and cell lysis conditions were identical to those used for T1-SpyTag. Turquoise-SpyC was purified by immobilized metal affinity chromatography using a nickel–nitrilotriacetic acid agarose gravity column (QIAGEN).

Protein purity was assessed by SDS–PAGE using 4–20% Mini-PROTEAN® precast gels (Bio-Rad). For SDS–PAGE analysis, any insoluble protein fractions were solubilized in 8 M urea prior to loading. Protein concentrations were determined using the Bradford assay (Bio-Rad).

### Initial formation of T1-SpyTag fibers and sheets

Following purification, T1-SpyTag was initially obtained as a soluble protein solution in elution buffer. During routine handling, spontaneous formation of fibrous structures was readily observed, with gentle agitation of the solution in microcentrifuge tubes leading to the appearance of visible fiber-like structures. Upon concentration using Amicon ultracentrifugal filters, sheet-like structures formed on the membrane surface of the concentrator. These sheet materials could be gently detached by pipetting, yielding macroscopically visible films. The resulting fibers and sheets were subsequently collected and used for downstream assembly, cargo-loading, and characterization experiments.

### Solution-state cargo loading

For solution-state cargo loading, pre-assembled T1-SpyTag fibers and sheets were collected by centrifugation at 8,000 × *g* and washed repeatedly with TBS buffer to remove unassembled protein. The T1-SpyTag macromolecular structures were then incubated with Turquoise-SpyC at 4 °C overnight to allow covalent conjugation via the SpyTag/SpyCatcher reaction.^15^ Unbound proteins were removed by three rounds of centrifugation and washing. The conjugated structures were resuspended in TBS and analyzed by optical and fluorescence microscopy. For verification of covalent cargo attachment, the assemblies were dissolved in 8 M urea and analyzed by SDS–PAGE.

### Dried-state cargo loading

For dried-state cargo-loading assays, 200 µL of T1-SpyTag solutions at concentrations of 0.375, 0.75, 1.5, and 3 mg mL^−1^ in elution buffer were deposited into 96-well plates. After air drying at room temperature, the wells were gently washed with deionized water and incubated with Turquoise-SpyC overnight at room temperature. Unbound Turquoise-SpyC was removed by at least five washing steps. Cargo conjugation onto dried T1-SpyTag sheets was confirmed by fluorescence microscopy and SDS–PAGE after dissolving the sheet layers in urea buffer.

### Assembly assay

Assembly of T1-SpyTag was monitored by turbidity measurements over a wavelength range of 300–450 nm. Aliquots (200 µL) of T1-SpyTag solutions at concentrations of 0.375, 0.75, 1.5, and 3 mg mL^−1^ in the corresponding buffer were loaded into 96-well plates. The apparent absorbance values, attributed to light scattering, were recorded at 10 min and 24 h using an Agilent BioTek Epoch 2 Microplate Spectrophotometer, with buffer-only samples serving as blank controls.

### Optical and fluorescence microscopy

Assembled T1-SpyTag fibers and sheets were imaged using an EVOS FL Color imaging system (Life Technologies) under transmitted light. Fluorescence images were acquired using the GFP filter cube to visualize Turquoise-SpyC conjugation.

### Transmission electron microscopy

For negative-staining TEM, 10 µL of T1-SpyTag solution was applied to a Formvar/carbon-coated 200-mesh copper grid and incubated for 1 min. The grid was washed twice with 10 µL of deionized water and blotted with filter paper. Subsequently, 10 µL of 1% (w/v) uranyl acetate solution was applied for 60 s before blotting nearly dry. Prepared grids were imaged using a JEOL JEM-1400 Flash transmission electron microscope operated at 100 kV.

### Atomic force microscopy

AFM imaging for air-dried sample was performed using an Asylum Research atomic force microscope operated in tapping mode. For sample preparation, 10 µL of T1-SpyTag solution (1 mg mL^−1^), prepared in either deionized water or TBS buffer, was deposited onto freshly cleaved mica substrates and incubated for 30 min. Samples were gently blotted with filter paper and dried under a stream of nitrogen gas prior to imaging. Silicon cantilevers suitable for tapping mode (AC240TS-R3, Oxford Instruments) were used. Images were acquired at room temperature with scan rates typically in the range of 0.2–0.5 Hz. AFM images were processed using standard flattening procedures. AFM imaging in solution was performed using an Asylum Research Cypher VRS system operated in tapping mode. Silicon wafers were first treated with UV/Ozone to clean the surface, followed by functionalization with a 1 mM Spermidine solution. Next, 10 µL of assembled T1-SpyTag fibers and sheets solution, stored in TBS buffer, was deposited onto the SiO2 substrate and incubated for 10 min. Samples were blow-dried under a stream of nitrogen gas, and rehydrated in TBS buffer prior to imaging. Silicon cantilevers suitable for tapping mode (Olympus AC40TS-R3) were used. Images were acquired at room temperature with scan rates typically in the range of 0.2–0.5 Hz. AFM images were processed using Asylum Research Offline Ergo software.

### Scanning electron microscopy

T1-SpyTag solutions were drop-cast onto silicon wafer substrates (Ted Pella, Inc.) and air-dried at room temperature overnight. The wafers were mounted on aluminum stubs using conductive epoxy and sputter-coated with a ∼2.7 nm layer of iridium using a turbo-pumped sputter coater (Q150T, Quorum Technologies) under an argon atmosphere. SEM imaging was performed using a field-emission scanning electron microscope (JEOL 7500F, JEOL Ltd.).

### Grazing-incidence Transmission Small-Angle X-ray Scattering (GTSAXS)

The T1-SpyTag was assembled into fibers and sheets in Tris-buffered saline (20 mM Tris–HCl, 250 mM NaCl) and the X-ray scattering characterization was conducted at Beamline 12-ID-B at the Advanced Photon Source of Argonne National Laboratory. Due to the limited amounts of assembled samples, conventional transmission small-angle X-ray scattering was unable to produce meaningful data. Instead, the sample was applied on silicon wafer surface, and analyzed with grazing incidence transmission small-angle X-ray scattering after drying. The X-ray incident angle was set at 0.1 degrees, which increased the X-ray footprint on the sample by approximately three orders of magnitude for thin films, thereby enhancing the X-ray scattering signals. Measurements were performed using an Eiger2 9M detector (DECTRIS LLC). The X-ray wavelength was set at 0.932 Å and the detector was positioned at 2 meters away from the sample. Calibration of the X-ray scattering vector (q) and conversion of the 2D GTSAXS image data into 1-D GTSAXS profile were carried out using the beamline software package matSAXS (https://12idb.xray.aps.anl.gov/Software_Processing.html). The X-ray scattering vector (q) is defined as q = 4πsin (θ)/λ, where λ represents the X-ray wavelength and 2θ corresponds to the Bragg angle.

### Disassembly assays

To evaluate the disassembly behavior of T1-SpyTag films, the macromolecular protein structures (100 µL, 4.4 mg mL^−1^) were deposited into 96-well plates and air-dried at room temperature. Film stability under different urea concentrations and pH conditions was assessed by monitoring changes in turbidity at 350 nm using a microplate spectrophotometer.

### Urea-induced disassembly

Air-dried T1-SpyTag films were first washed three times with deionized water. Subsequently, 100 µL of deionized water was added to each well, and the corresponding absorbance at 350 nm was recorded as the initial water control (A_H2O_). The water was then removed and replaced with 100 µL of urea solutions at final concentrations of 0, 2, 4, 6, or 8 M. Turbidity was monitored at room temperature over a wavelength range of 300–450 nm, with the apparent absorbance at 350 nm recorded every 20 min for 3 h. The absorbance of empty wells was used as background control (A_empty_). The disassembly was monitored as: Assembly ratio (%) = 1- (A−A_empty_)/(A_H2O_−A_empty_)×100, where *A* represents the apparent absorbance at a given time point.

### pH-induced disassembly

Air-dried T1-SpyTag films were rehydrated with Tris-buffered saline (20 mM Tris–HCl, 150 mM NaCl) adjusted to pH 6.0, 7.6, or 8.5. The apparent absorbance at 350 nm was recorded immediately after buffer addition and defined as the initial absorbance (A_0_). Samples were gently agitated every 20 min, and turbidity was monitored over a total duration of 3 h. The absorbance of empty wells was used as background control (A_empty_). The disassembly was monitored as: Assembly ratio (%) = 1- (A−A_empty_)/(A_0_−A_empty_)×100, where *A* represents the apparent absorbance at a given time point.

### Statistical analysis and visualization

All experiments were performed in at least triplicate unless otherwise stated. Data are presented as mean ± standard deviation (SD). Statistical analyses were performed using GraphPad Prism 9.0 (San Diego, CA, USA). Molecular visualization and figure rendering were performed using PyMOL 3.10. Three-dimensional conceptional schematics were made in Blender 3.6.

## Supporting Information

Supporting Information is available from the Wiley Online Library or from the author.

## Acknowledgements

This research was supported as part of the Center for Catalysis in Biomimetic Confinement (CCBC), an Energy Frontier Research Center funded by the U.S. Department of Energy (DOE), Office of Science, Basic Energy Sciences (BES). At Michigan State University, the work was supported under award DE-SC0023395, at Argonne National Laboratory under contract DE-AC02-06CH11357, and at the Molecular Foundry at Lawrence Berkeley National Laboratory under contract DE-AC02-05CH11231. A portion of this research was performed using Advanced Photon Source (APS) beam time award(s) (DOI: https://doi.org/10.46936/APS-191708/60015433) from the APS, a DOE Office of Science user facility at Argonne National Laboratory, and is based on research supported by the DOE Office of Science-BES, under Contract No. DE-AC02-06CH11357. We thank Markus Sutter and Cheryl Kerfeld for plasmid constructs, Morgen Clark for initial AFM data collection, Megan Cross for cryo-electron microscopy screening, and staff at the MSU Center for Advanced Microscopy for TEM and SEM assistance.

## Author Contributions

Y.W. conceived the project and performed protein purification, assembly, disassembly, cargo-loading assays, UV–visible spectroscopy, transmitted-light microscopy, fluorescence microscopy, TEM, and SEM experiments. Y.W. also analyzed the data, wrote the original draft, and prepared all figures. X.Z. performed GT-SAXS experiments and its data analysis. Y.W. and P.D.A. performed AFM imaging and data analysis. Y.W. X.Z., Y.W., and P.D.A. contributed to manuscript revision. R.P.H. supervised the project, provided funding support, and edited the manuscript.

## Data Availability Statement

The data that support the findings of this study are available from the corresponding author upon reasonable request.

