## Supplemental Information for "SpyTag-Enabled Assembly of Bacterial Microcompartment Trimers into Macroscopic Layered Protein Materials"

**T1-SpyTag sequence (SpyTag is in red):**
MHHHHHHMDHAPERFDATPPAGEPDRPALGVLELTSIARGITVADAALKRAPSLLLMSRP
VSSGKHLLMMRGQVAEVEESMIAAREIAGAGGGSGGSAHIVMVDAYKPTKGGSGGSGA
LLDELELPYAHEQLWRFLDAPVVADAWEEDTESVIIVETATVCAAIDSADAALKTAPVVL
RDMRLAIGIAGKAFFTLTGELADVEAAAEVVRERCGARLLELACIARPVDELRGRLFF*

**Turquoise-SpyCatcher sequence (SpyCatcher is in red):**

MSYYHHHHHHDYDIPTTENLYFQGAMVDTLSGLSSEQGQSGDMTIEEDSATHIKFSKRD
EDGKELAGATMELRDSSGKTISTWISDGQVKDFYLYPGKYTFVETAAPDGYEVATAITFT
VNEQGQVTVNGKATKGDAHIGGGSGGASVSKGEELFTGVVPILVELDGDVNGHKFSVS
GEGEGDATYGKLTLKFICTTGKLPVPWPTLVTTLSWGVQCFARYPDHMKQHDFFKSAM
PEGYVQERTIFFKDDGNYKTRAEVKFEGDTLVNRIELKGIDFKEDGNILGHKLEYNYFSD
NVYITADKQKNGIKANFKIRHNIEDGGVQLADHYQQNTPIGDGPVLLPDNHYLSTQSKL
SKDPNEKRDHMVLLEFVTAAGITLGMDELYK*

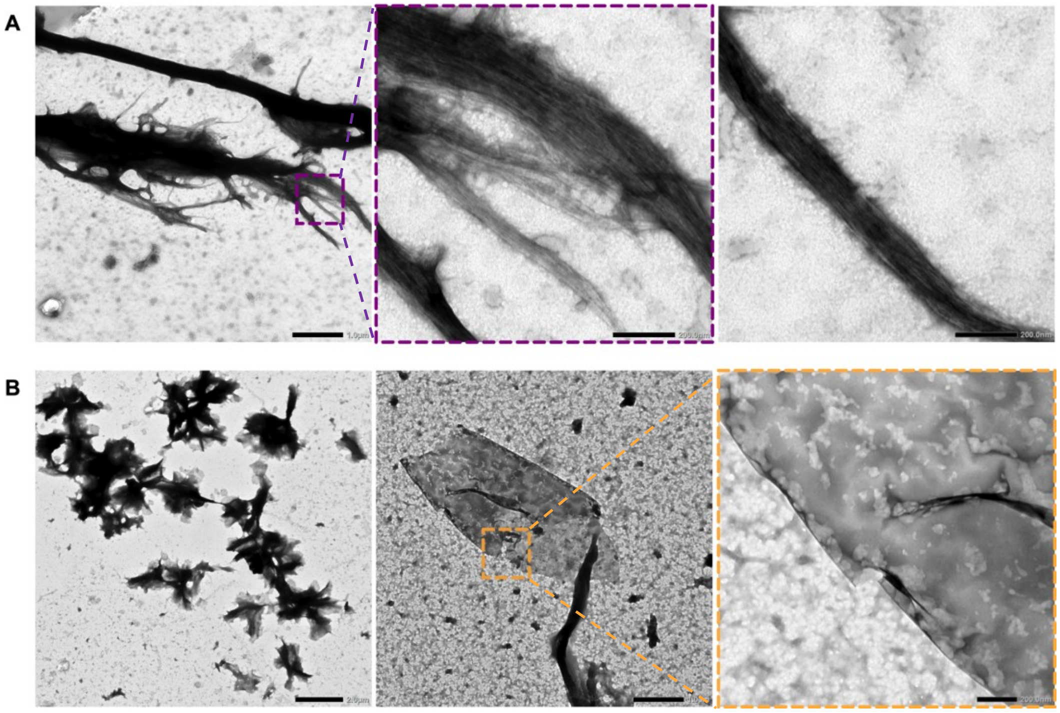


**Figure S1.** Transmission electron microscopy (TEM) images of assembled T1–SpyTag structures. **(A)** Representative TEM images showing fiber-like assemblies formed by T1–SpyTag after purification and incubation. Higher-magnification views (dashed boxes) reveal bundled filamentous morphology and internal alignment features along the fiber axis. **(B)** Representative TEM images of sheet-like T1–SpyTag assemblies displaying extended, plate-like morphologies. Boxed regions indicate areas selected for higher-magnification views, showing textured and multilayered appearance at the sheet surface. All samples were negatively stained. Scale bars are as indicated.

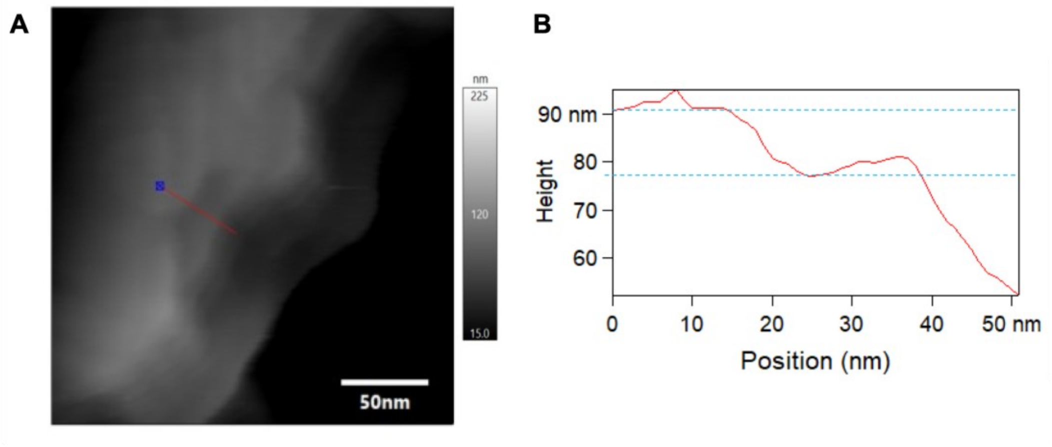


**Figure S2.** Representative AFM height image of T1-SpyTag sheets in TBS solution. (A) AFM images of T1-SpyTag sheets. (B) Height profile from the red transit in (A).

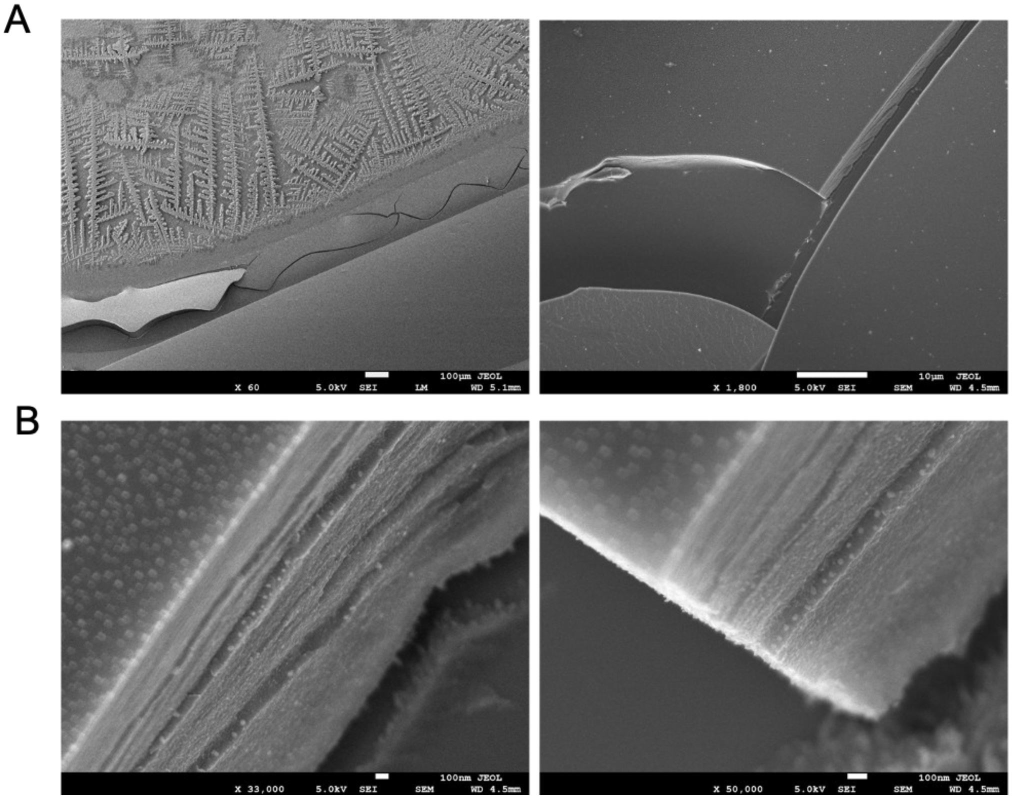


**Figure S3.** Scanning electron microscopy (SEM) images of air-dried T1–SpyTag materials showing multilayered morphology and interlayer separation features. **(A)** Low- and **(B)** high-magnification views reveal stacked, sheet-like architectures with visible layer boundaries and crack-like gaps between adjacent layers in several regions.
